# Early bird dynamics reduce selection imposed by species interactions in boom-and-bust regimes

**DOI:** 10.64898/2026.10.26.761801

**Authors:** Nittay Meroz, Jonathan Friedman

## Abstract

Interspecific interactions are a major force shaping evolution, yet the magnitude of their effect varies across ecosystems and interactions. What explains this variability remains underexplored. Here, we propose that the temporal dynamics of growth and resources are a key determinant. We study how the degree to which species interactions alter selection forces depends on a system’s growth regime, ranging from steady chemostat-like growth to boom-and-bust cycles. Using consumer-resource simulations together with analytical approximations, we show that selection in boom-and-bust cycles is dominated by an *early bird* effect: because most generations elapse early in a growth cycle — when populations are small and resources are abundant — selection is concentrated in a period where species interactions are relatively weak. As a result, boom-and-bust dynamics diminish the degree to which growth rate reducing interactions alter selection. In contrast, interactions that act through mortality, such as the secretion of lethal toxins, operate regardless of growth phase and are therefore not reduced, and can even be amplified in boom-and-bust dynamics. These findings help contextualize estimates of the evolutionary effects of species interactions across experimental systems, and suggest that the growth regime is a coarse-grained, measurable feature that governs when biotic interactions leave a strong evolutionary mark.

## Introduction

From finches adapting their beak size to the presence of other species, to bacteria diverging in niche use during co-evolution, or plants synthesizing toxins to deter herbivores, the evolutionary effects of species on each other are apparent throughout the tree of life and have been recognized since the early days of the theory of evolution [1, 2, 3]. However, the extent to which species evolution is shaped by biotic versus abiotic factors is of long-standing debate [4]. Understanding when biotic interactions generate pronounced evolutionary effects is crucial for explaining past evolutionary trajectories and forecasting future evolutionary dynamics [5]. A general resolution for this debate seems unfeasible, as the relative importance of biotic and abiotic factors likely differs substantially across ecological systems. Nevertheless, a path forward may lie in identifying general features that tip the balance towards one or the other.

Recent empirical studies provide measurements of species evolutionary effects on each other, either by evolving species with and without a specific interaction [6, 7, 8, 9, 10], or by measuring the fitness of mutants in the presence and absence of another species [11, 12, 13, 14]. Indeed, these studies demonstrate that interspecific interactions can reshape species’ adaptive landscapes; yet, they reveal substantial variation in the magnitude of this effect [15]. The correlation between the fitness of mutants in the presence vs. in the absence of an interaction ranges from 0.98— indicating that the interaction barely alters selection in some studies [13] to 0.6 in others, where its effect is substantial [12]. Why do species leave a clear evolutionary mark on each other in some cases and only a faint one in others? Naively, one might expect that stronger ecological interactions — where species affect each other’s population sizes substantially— should produce stronger evolutionary effects. However, this expectation is not grounded in theory and does not hold empirically — strongly interacting species often affect each others’ evolution only weakly [6], while seemingly weak interactions could have a strong effect on evolution [10]. While recent theoretical advances have made progress in formalizing the mapping from ecological interactions to selection [15], we still lack a general understanding of the factors affecting the extent to which interactions shape selection and the manner in which they do so. Here, we show that one key factor is whether populations experience steady growth or undergo boom-and-bust cycles driven by fluctuating resource availability and mortality.

Fluctuating conditions are common both in natural and laboratory settings. Natural populations are often far from a steady state; nutrient inflows, weather disturbances, or pathogen invasions can lead to rapid growth or a swift decline [16]. Different ecosystems can be characterized by the degree of disturbance they experience, ranging from relatively steady environments with balanced growth and death to highly dynamic ones in which populations often undergo rapid expansions and collapses — sometimes referred to as boom-and-bust dynamics [17, 18]. For example, phototrophs in the oligotrophic open ocean have a fairly constant population size [19]; while individual bacterial taxa in coastal waters, where nutrient pulses are more frequent, can increase or decline 20-fold in abundance over the course of a single week [20]. Notably, most experimental evolution studies with bacteria also occupy the boom-and-bust end of this spectrum, as serial dilution protocols impose cyclic rounds of growth and resource depletion [21]. The degree to which an environment is characterized by steady turnover or by boom-and-bust dynamics is not merely a background feature of ecosystems; it has implications for their stability, composition, and for which traits are under selection [22, 23, 24].

Theoretical studies of evolution in steady, chemostat-like conditions often find strong evolutionary consequences of species interactions [25, 26, 27, 28]. However, the evolutionary consequences of interaction in boom-and-bust conditions remain poorly understood. Notably, boom-and-bust dynamics are distinct from general fluctuations in environmental parameters, such as temperature, in that an externally driven disruption, such as a nutrient pulse or dilution event, is followed by environmental changes driven by the populations’ own rapid growth. Here, we use mathematical modeling and numerical simulations to ask: how do the effects of interspecific interactions on evolution depend on the growth dynamics of the system, ranging from steady, chemostat-like conditions to fluctuating, boom-and-bust cycles? We model environment-mediated species interactions, as is common for bacteria [29], focusing on resource competition and toxin secretion, and derive general results applicable to other environment-mediated mechanisms. We find that when species interact by inhibiting each others’ growth (e.g., via resource competition), boom-and-bust dynamics give rise to ‘early bird’ effects[22]: the largest fitness gains come from increasing growth rates when resources are abundant and populations are small — a phase in which interspecific interactions are weakest. As a result, boom-and-bust dynamics tend to reduce the evolutionary consequences of growth-related interactions. However, when species interact by affecting each others’ death rates (e.g., secreting a toxin that kills the other[30]), a more complex picture emerges where boom-and-bust dynamics could, in some cases, amplify the effect of the interaction. These results help interpret findings from experimental evolution studies and provide testable predictions across natural ecological contexts.

## Results

### Simulating resource competition across growth regimes

We employ a classic consumer-resource model in which two species compete for two substitutable resources (Fig. 1A, Section S1). Briefly, we model each species’ growth rate as the sum of the consumption rates of available resources. Species *i* consumes resource *j* at a maximum rate *c*_*ij*_ modulated by a consumption function *f*_*ij*_(*R*_*j*_) that depends on resource concentration with a Monod-type saturating form. Both population and resource concentrations are diluted at a rate *d*(*t*), and new resources flow-in at the same rate *d*(*t*) with a concentration of *R*_*j,in*_.

**Figure 1:**
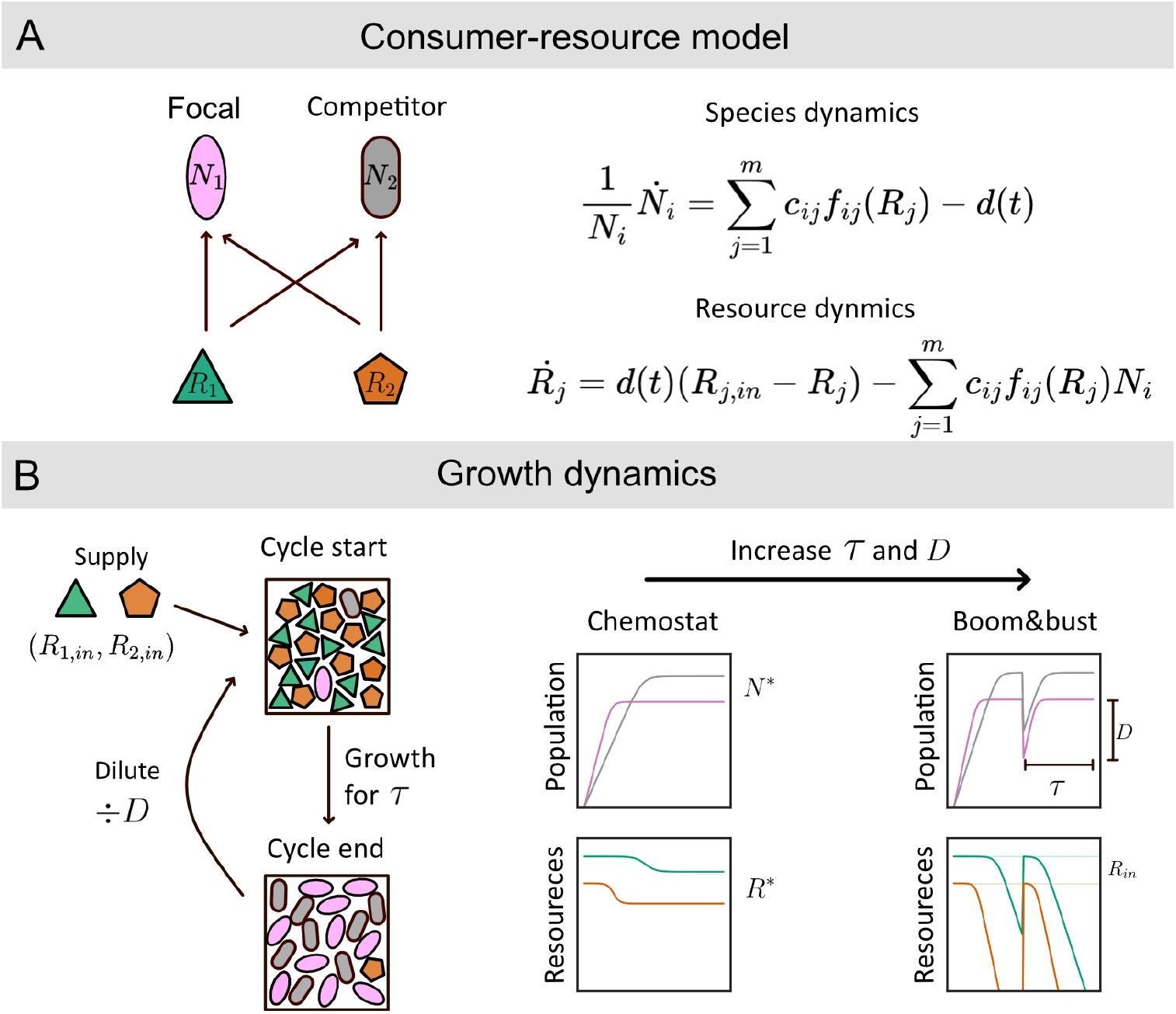
Modeling resource competition across growth regimes. (A) Two species compete for two substitutable resources. Population growth x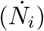 and resource depletion 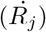 are governed by speciesspecific consumption rates, which are given by the max rate *c*_*ij*_ and the concentration-dependent function *f*_*ij*_(*R*_*j*_). In the chemostat-limit the system experiences a washout rate *d*, while fresh resources enter at concentrations *R*_*j,in*_. (B) Growth conditions vary along a continuum from continuous flow (chemostat) to discrete growth-dilution cycles (boom-and-bust). Chemostat conditions maintain a constant flow rate *d*. Boom-and-bust conditions involve cycles of batch growth for duration *τ* followed by an instantaneous dilution event by factor *D*. The time-averaged flow rate is kept constant across all regimes by setting *D* = *e*^*τd*^ (Section S2.2)

We vary the growth regime between chemostat-like conditions and boom-and-bust cycles by varying the temporal pattern of the flow rate *d*(*t*) (Fig. 1B, Section S2). In a chemostat, flow is continuous and constant: 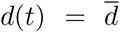. We depart from chemostat dynamics by discretizing the flow into dilution-growth cycles, in which discrete dilute-and-replenish events are separated by growth periods of length *τ* during which there is no flow [31]. We define *D* as the discrete dilution factor applied every *τ*. To enable comparison between regimes, we match their time-averaged flow rates by imposing the scaling relation 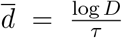, which also equalizes the mean nutrient supply (Section S2.2). In this manner, when *τ* is small, the variability in growth and death rates within a cycle is small, and the dilutions become smaller (approaching 1 as the cycle length approaches 0). As *τ* increases, populations experience increasingly pronounced boom-and-bust dynamics, and *D* increases.

### Selection differs between chemostat and boom-and-bust dynamics

First, we ask whether different growth regimes impose different selection. To do so, we introduce mutants into an environment where only their ancestor is present (monoculture) and quantify their invasion fitness over one growth cycle (Eq 7). A mutant inherits all of its ancestor’s parameters with a small, randomly drawn change that can be either positive or negative for each parameter (Methods).

As previously established [31], the fitness of a mutant is not affected by the growth regime if the environment contains only a single resource type and consumption is a linear function of resource concentration (Fig. S1). However, this equivalence breaks down if consumption is nonlinear, or if there are multiple resources, even if each is consumed linearly (Fig. S2A). The reason is that while the consumption of each individual resource remains linear, the total growth rate over the time cycle depends nonadditively on the temporal variation of the resources [31, 32]. This is due to the fact that the concentration of the resources is correlated, as their consumption depends on population size (Section S3.3).

### Boom-and-bust dynamics reduce the affect of species interaction on selection

The presence of another species modulates selection by altering resource concentrations. This can be seen by comparing the fitness of mutants when only their ancestor is present (monoculture) to the fitness of the same mutants in the presence of both their ancestor and a competitor (coculture). Mutant fitness can differ substantially between these two conditions, both in magnitude and in sign, a mutation that is beneficial in monoculture can be detrimental in coculture, and vise versa (Fig 2A). Here, we do not assume a specific tradeoff between the utilization rates of each resource; but sample each parameter independently (Table S1). Previous work has established that, depending on the shape of the tradeoff curve, such differences in selection would lead to either niche differentiation or niche convergence [33, 22, 26].

**Figure 2:**
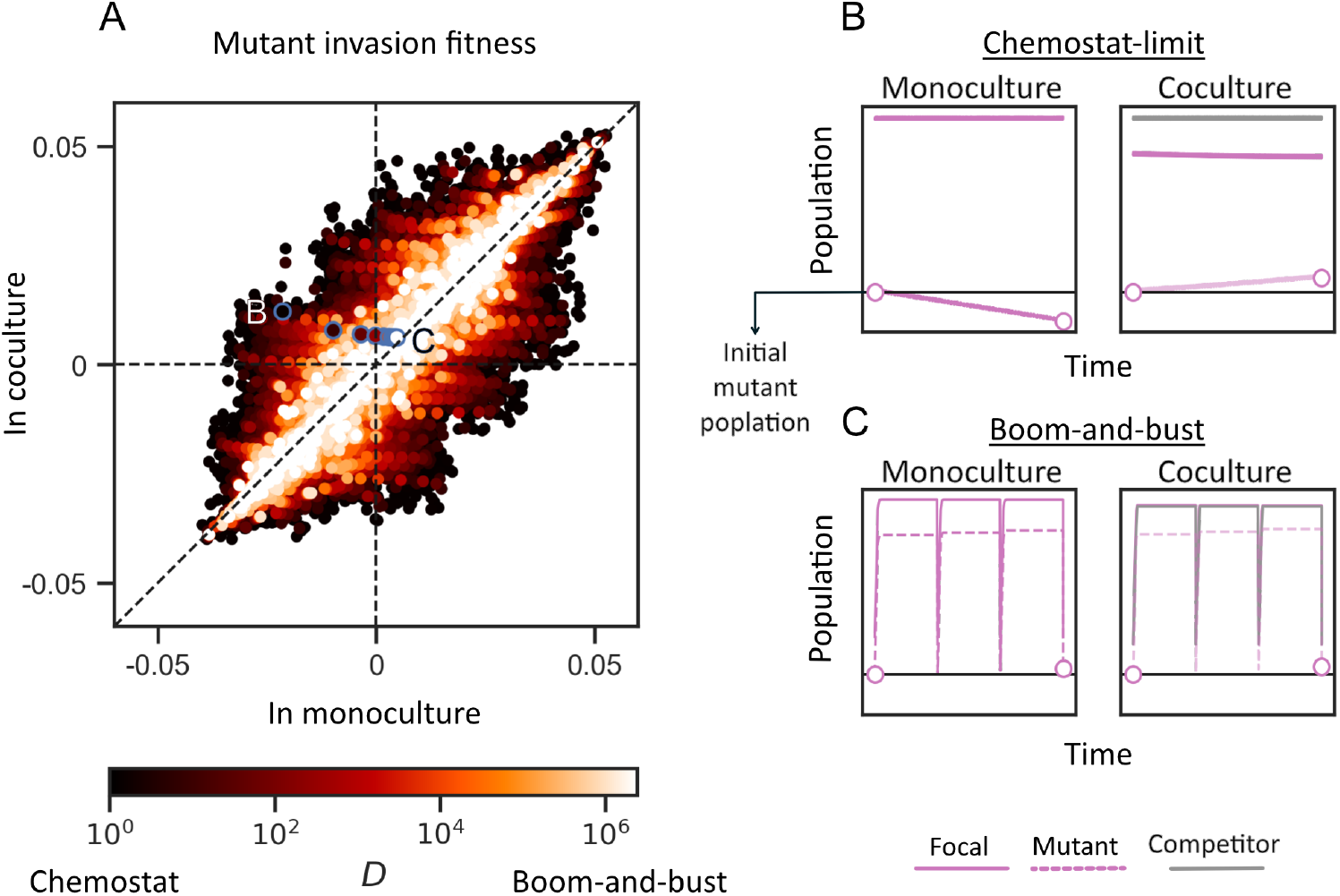
Evolutionary effects of resource competition are reduced in a boom-and-bust regime. (A) Mutant invasion fitness (Eq. Eq 7) at steady state in coculture versus monoculture (mutant invades to a culture where only its ancestor is present). Points represent 30 mutations across 4,607 unique resource competition simulations (after filtering, see Methods and Fig.S3) each evaluated under 9 *τ* values. Full parameter list is found in Table S1. Point color denotes the strength of the boom-and-bust dynamics, mapped to the log dilution factor *D*. Blue circles trace a single mutant invading the same ancestral background across varying growth regimes. Panels A and B trace the population dynamics at the two extremes: in the chemostat limit (B), the mutant exhibits negative fitness in monoculture but positive fitness in coculture; conversely, under strong boom-and-bust dynamics (C), the mutant exhibits positive fitness in both environments.

Differences in selection that are driven by resource competition are substantially reduced in boom-and-bust dynamics (Fig. 2). When populations undergo weak dilution-growth cycles (dark dots, small *D*), the fitness of the mutant can vary substantially between monoculture and coculture. However, as the dilution factor *D* increases, these differences are reduced, and selection becomes relatively unaffected by the presence of the other species (white dots, approaching the 1:1 line). This reduction is not only an aggregated pattern but applies per-mutation; the blue circles depict a specific mutation in the same background coculture and the same average flow rate 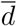, but under different growth regimes (different *D* and *τ*). In the chemostat limit, this mutation has contrasting effects in monoculture and coculture (Fig. 2B: negative in monoculture, positive in coculture), whereas the difference disappears in the boom-and-bust limit (Fig. 2C: selection is positive in both mono- and coculture).

### Selection in boom-and-bust dynamics is strongest for growth early in the cycle

To explain why boom-and-bust dynamics reduce the evolutionary effect of species interactions, we consider how the environment changes over the course of a single cycle. Because growth is multiplicative, the fitness of a mutant in a time-varying environment is determined by its geometric mean growth rate [34]. Therefore, growth ability in conditions experienced over more generations contributes more strongly to overall fitness. When the environment changes due to the metabolic activity of the populations themselves, the rate of environmental change is coupled to population size. Early in each cycle of a typical boom-and-bust scenario, population sizes are small relative to the biomass that the supplied resources can support (Fig. 3A). Therefore, resource depletion is slow, and resource concentrations remain close to their initial values over many generations. As populations grow, resource depletion accelerates until resources are exhausted. Because more generations elapse during the early resource-rich phase, selection is disproportionately concentrated in this window. Following analogous usage in an ecological context, we refer to this bias toward early, resource-rich conditions as the early bird effect [22, 35].

**Figure 3:**
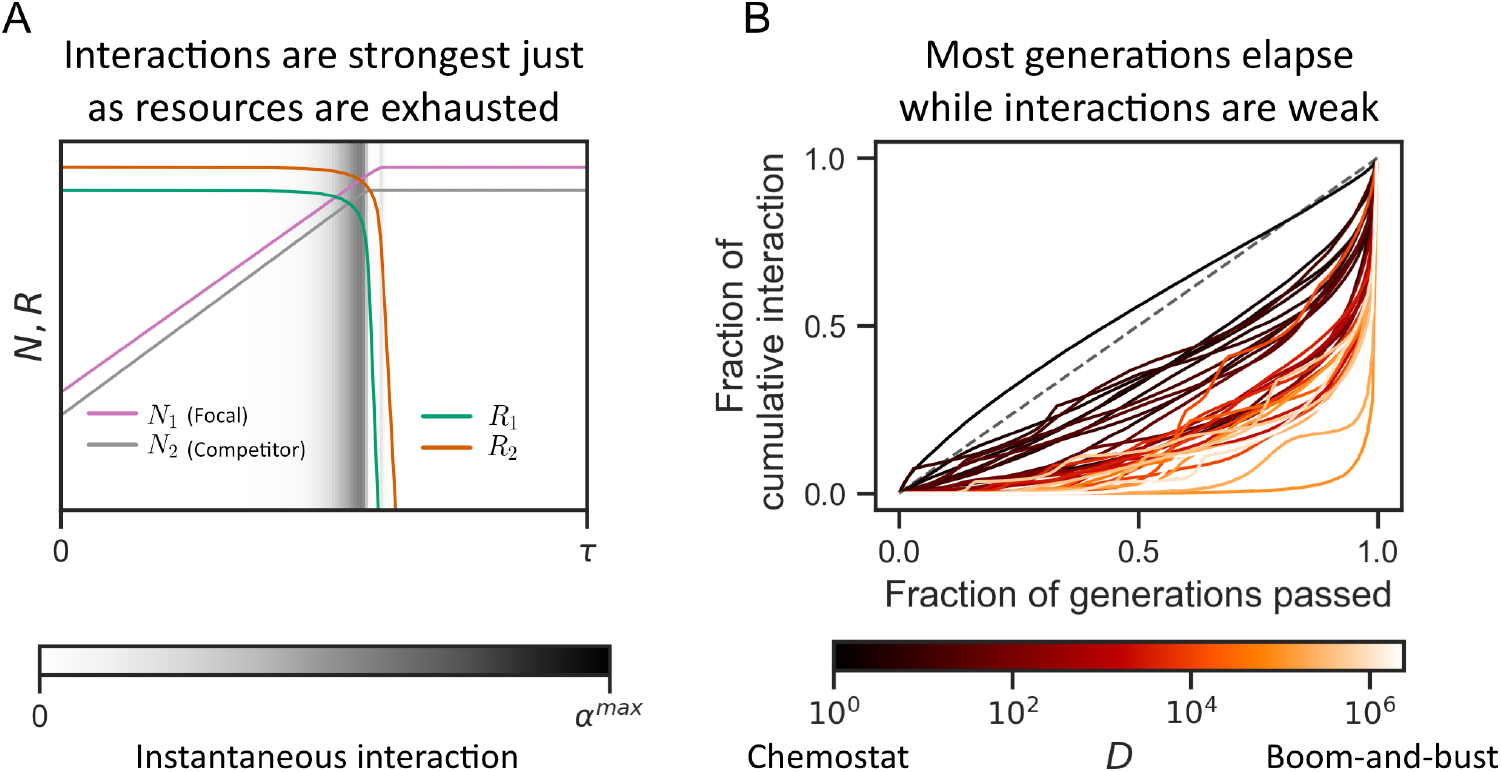
In boom-and-bust cycles, species interactions are weak for the majority of the cycle and peak before resources are exhausted. (A) Representative population (*N*) and resource (*R*) trajectories for two species competing for resources *R*_1_ and *R*_2_. The grayscale background shading indicates the instantaneous strength of competition imposed by the competitor on the focal species. (B) The cumulative fraction of total interaction is plotted against the cumulative fraction of generations passed within a cycle. The dashed diagonal line represents a hypothetical uniform interaction rate per generation. The pronounced rightward bow of the curves demonstrates that interactions are heavily concentrated in the final few generations of the growth cycle. Line color maps to the dilution factor log(*D*), representing data binned across 67 *D* values. The plot includes data from 4, 607 simulations, parameter list is found in Table S1.

Because species interact through resource competition, the rates at which resources are depleted determine the strength and timing of interspecific interactions. In a typical boom-and-bust cycle, the total instantaneous interaction [36](Methods), which quantifies the rate at which species alter each other environment in a growth relevant manner, is negligible for most of the cycle and peaks briefly just before resources are exhausted (Fig. 3A). To quantify how much of the focal species’ growth is affected by this interaction, we plot the fraction of cumulative interaction against the fraction of generations elapsed (Fig. 3B), as a proxy for the potential effect of the interaction on selection. When the growth regime is close to a chemostat (low *D*), species interact throughout the cycle, as both population and resource densities remain near equilibrium (dark lines, close to the 1:1 line). However, under strong boom-and-bust cycles, species interact mostly in the last few divisions (light lines), right before resources are exhausted.

### Selection in boom-and-bust cycles is dominated by growth on input resource concentrations

In resource competition, growth rates, and thus selection, are determined by resource concentrations. In a chemostat, species experience constant, steady-state resource concentrations (*R*^*\**^), which depend on species consumption rates of the different resources. Differences in selection between monoculture and coculture ultimately reflect the way the additional species modify *R*^*\**^ (Section S3.3). However, in boom-and-bust cycles, species experience fluctuating resource concentrations, and thus different selection compared to chemostat conditions. These differences can be substantial even under mild boom-and-bust cycles (Fig. 4A).

**Figure 4:**
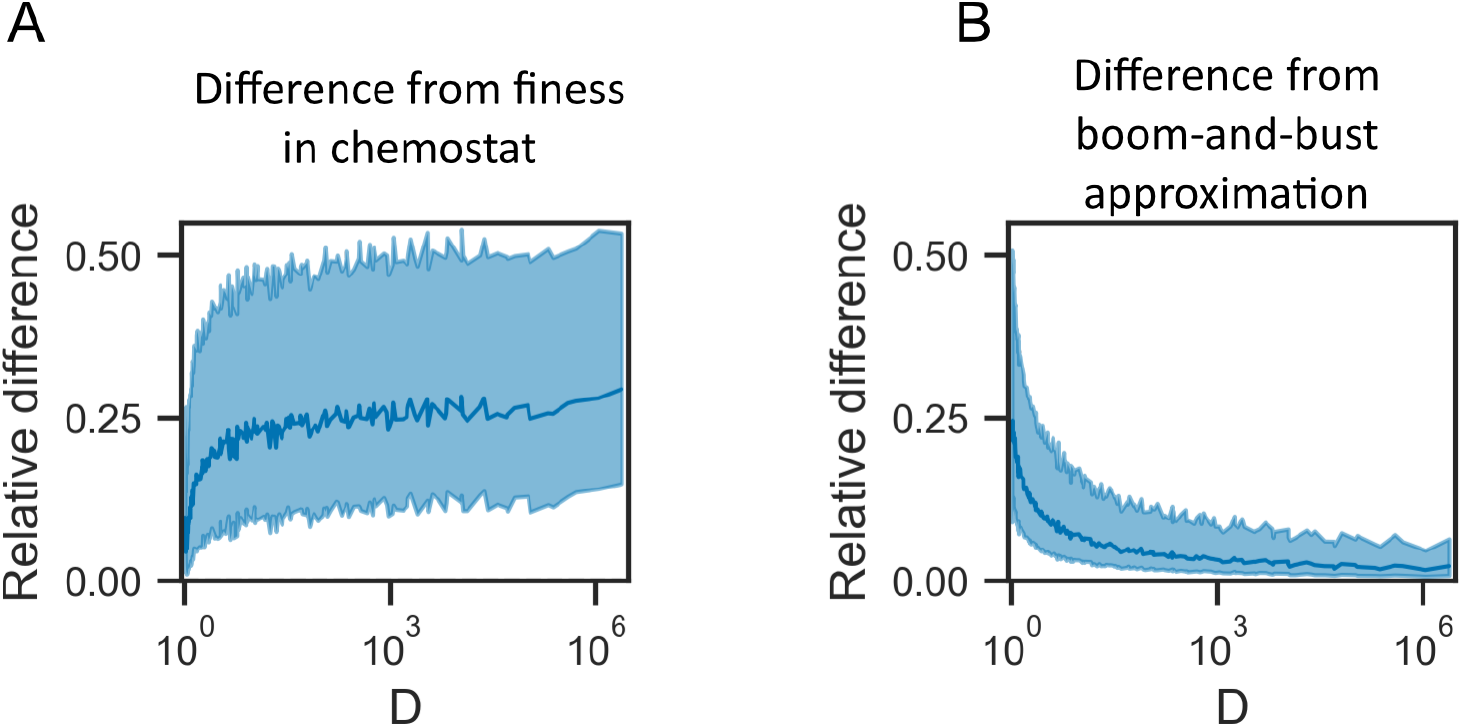
Selection in the boom-and-bust regime is dominated by external resource supply levels, independent of species presence. (A) The relative difference in mutant invasion fitness between the chemostat analytical solution (Eq S26) and simulations across varying dilution factors. (B) The relative difference in mutant invasion fitness between the boom-and-bust approximation (Eq S27) and simulations. For panels, solid lines and shaded regions represent the mean and inter-quartile range, respectively. The relative difference is defined as (*S*_*sim*_*™S*_*ana*_)*/*(*S*_*sim*_+*S*_*ana*_), where *S*_*sim*_ is the simulated fitness and *S*_*ana*_ is the analytical approximation. Table S1 lists the parameters used to produce these simulations.

In strong boom-and-bust regimes, selection is dominated by species’ growth rate on input resource concentrations, which is unaffected by the presence of other species (Section S3.3). In this regime, populations typically have enough time to completely exhaust resources by the end of the cycle; as a result, resource concentrations at the start of each cycle are determined solely by external supply. As established earlier, resource depletion accelerates due to exponential bacterial growth, and most resources are consumed in a brief window near the end of the growth within the cycle. Taking this to the limit, we can approximate resource concentrations as a step function — held at the input concentration until abruptly exhausted. Under this approximation, mutant fitness reduces to its growth rate on input resource concentration relative to that of the resident (see Section S3.3 for the complete approximation). This approximation captures mutant fitness well under boom-and-bust dynamics (Fig. 4B) and explains why species interactions matter less in this regime: species alter each other’s environment through resource depletion, but in the boom-and-bust limit, selection occurs mostly before substantial depletion has taken place.

### Interactions that effect species death rates are under strong selection even in boom-and-bust regime

To test whether these conclusions extend beyond resource competition, we consider species that interact solely through the secretion of a toxin (Fig. 5A, Section S6). In this scenario, each species grows on a separate resource, and one species secretes a toxin as it grows. The toxin can either inhibit the growth rate of the focal species (a bacteriostatic toxin) or kill it at some rate (a bactericidal toxin). Both bacteriostatic and bactericidal toxins affect selection through two mechanisms. First, they directly alter the growth or death rate of the focal species; a mutant that is less sensitive to the toxin has a fitness advantage. Second, by reducing the affected species’ population size, the toxin indirectly increases the concentration of the resources that that species’ consumes, and thus affects selection on traits related to resource consumption (Section. S6.1). As a result, a mutation can be beneficial in monoculture but detrimental in coculture, or vice versa if there is a tradeoff between maximal growth rate and toxin susceptibility.

**Figure 5:**
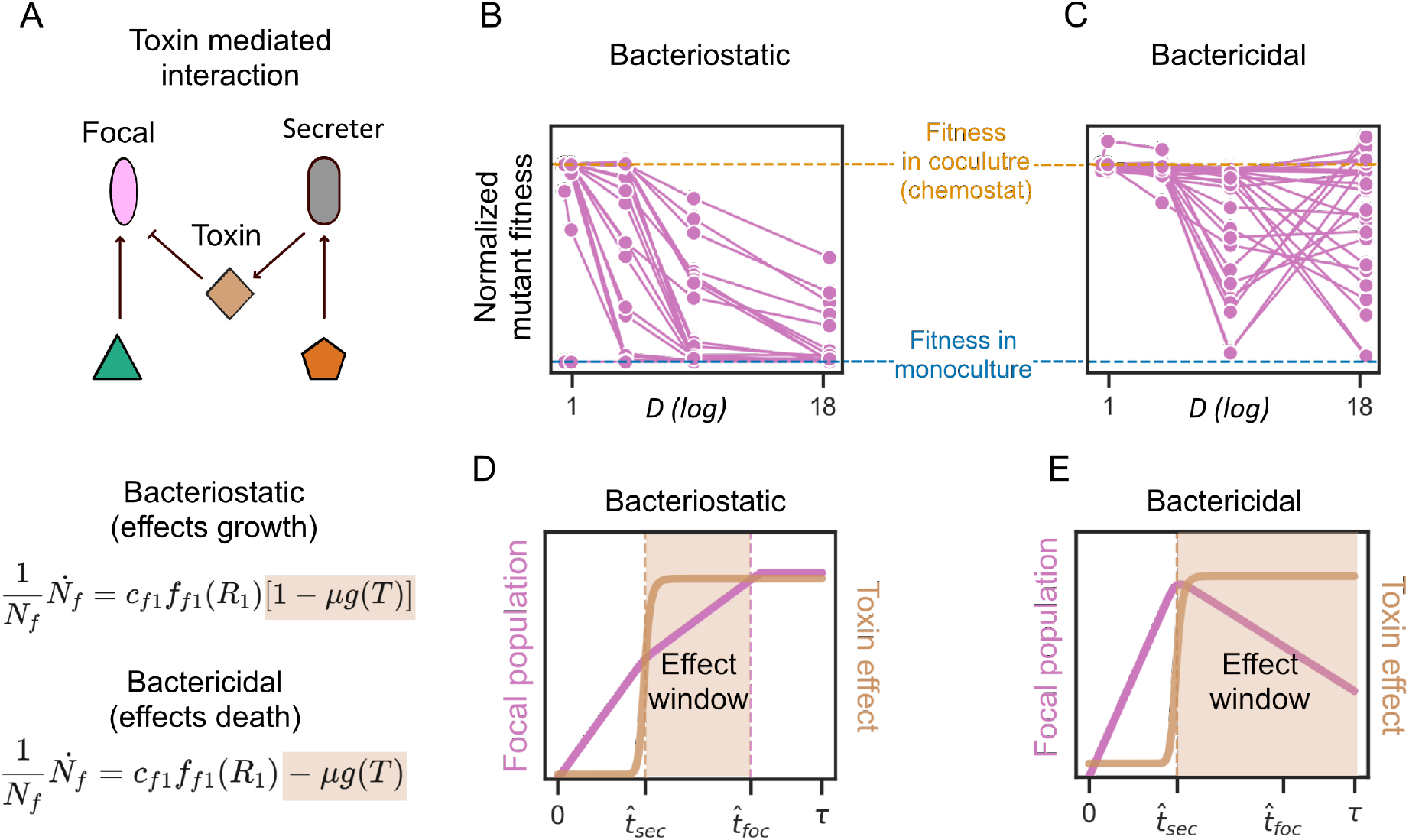
Bacteriostatic and bactericidal toxins differ in how boom-and-bust dynamics modulate their evolutionary impact. (A) Two species interact solely through the secretion of a toxin. We ask how much the evolution of the affected (focal) species is influenced by this interaction across growth regimes. We model two types of toxins: bacteriostatic toxins, which reduce the focal species’ growth rate, and bactericidal toxins, which increase its death rate. (B) Normalized fitness of a mutation conferring a 20% increase in growth rate and a 50% decrease in toxin resistance, plotted across dilution factors *D*, for a bacteriostatic toxin. Each dot represents a different background pair of secretor and focal strain; lines connect the same pair across *D* values (30 pairs shown). Fitness is normalized such that 1 corresponds to the mutant’s fitness in coculture under chemostat conditions *S*_*co*_ and 0 corresponds to its fitness in monoculture *S*_*m*_: |(*S*_*sim*_ *™S*_*m*_)*/*(*S*_*co*_ *™S*_*m*_)|. (C) Same as (B), but for a bactericidal toxin. Note that fitness can diverge from the monoculture value as *D* increases, indicating that boom-and-bust dynamics can amplify the evolutionary effect of bactericidal interactions. (D) Population size of the focal species (pink) and the bacteriostatic toxin effect *µg*(*T*) (orange) during an example growth cycle. The shaded region marks the interaction window — the period during which toxin levels are high and the focal species is still actively growing. (E) Same as (D), but for a bactericidal toxin. The shaded region now extends to the end of the cycle, reflecting that killing acts on cells regardless of growth.. Table S3 lists the parameters used to produce these simulations

To illustrate how these effects change with growth regime, we follow the fitness of a trade-off mutation, with a 20% increase in growth rate and a 50% decrease in toxin resistance across different background pairs and growth regimes. For bacteriostatic secretions, fitness converges consistently toward the monoculture value as *D* increases (Fig. 5B, approaching the blue line), similar to the pattern observed for resource competition (Fig. 2). However, for bactericidal secretions, we observe that fitness can either converge or diverge from the monoculture value as *D* increases (Fig. 5C) — meaning that boom-and-bust dynamics can, in some cases, amplify the evolutionary effect of species interactions.

To understand the difference between bacteriostatic and bactericidal toxins, we consider the temporal dynamics of toxin accumulation. Because the toxin is produced by growing cells, toxin levels are low at the beginning of the cycle, when the secretor population is small, and rise rapidly during exponential growth. The peak toxin concentration in a boom-and-bust cycle, necessarily exceeds the steady-state level in the equivalent chemostat while the concentration at the beginning of the cycle is lower (Section S6.1). Therefore, while in boom-and-bust dynamics the toxin has a stronger effect during part of the cycle, it has a weaker effect for the rest. Understanding whether boom-and-bust dynamics would reduce or amplify the effect of toxin related interactions can be boiled down to the relationship between how much the peak effect increases and the duration of the effect window.

The key difference between bacteriostatic and bactericidal interactions is in the duration of the effect window. For both toxin types, toxin concentrations exceed those of the chemostat when the secretor has grown substantially and thus depend on its growth rate. However, while a bacteriostatic toxin affects only until the focal population resumes growth (Fig. 5D), a bactericidal toxin continues to affect until the end of the cycle, regardless of the focal growth (Fig. 5E). The effect window for a bacteriostatic toxin depends on the difference in growth rates between the secreting and the focal species; if the secreting species grows slower than the focal one, it might not have any effect at all, and only when it grows substantially faster can a large fraction of the focal growth be affected. Such constraints do not apply to the bactericidal toxin—the effect window can be large even when the secretor grows substantially slower if *τ* is large. To formalize this logic and to check that it coheres with simulations, we developed approximations that simplify toxin and population dynamics into step-wise functions and therefore give rise to strict effect-windows (Sections S6.1, S6.2, Table S2) These approximations capture simulation results well under a wide parameter range (Fig. S6).

Finally, if peak toxin concentrations are substantially higher in boom-and-bust, this could counteract the narrowing of the effect window; however, this is unlikely to occur in the bacteriostatic case (Section S6.1). The reason is that the difference between chemostat and boom-and-bust concentrations is linked to the secretor growth rate compared to the flow rate (*d*); a fast growing species can almost completely exhaust its resources in a chemostat, reaching a population size similar to that of the boom-and-bust stationary phase—and producing the matching toxin concentrations. Only when the sector grows near washout — that is, barely growing faster than the dilution rate — does a substantial difference in toxin concentrations between chemostat and boom-and-bust arise (Fig. S4). However, this entails that the effect window is short—in the case that the secretor still grows slightly faster than the focal—or vanishes completely otherwise.

## Discussion

In this study, we demonstrate that the effect of growth-related species interactions on natural selection is expected to be reduced under boom-and-bust dynamics due to an ‘early bird effect’. Selection in a fluctuating environment does not necessarily equate to selection under the equivalent mean environment (the equivalent chemostat) due to nonlinearities and nonadditivity [37], and thus should be quantified as the geometric mean fitness across a changing environment [34]. However, here we show that in boom-and-bust dynamics—in which rapid growth drives environmental change—fitness gains are biased towards early growth because most generations elapse while cell densities, and hence interactions, are still low. These results are consistent with other studies that found selection for early growth within batch dynamics [35, 24] and with classic ecological theory, in which disturbed, resource-rich early-successional conditions favor fast-growing pioneer strategies over competitive ones [38, 39]. Our work departs from these by formalizing and addressing what this implies for the evolutionary effects species have on each other and how this differs between growth-driven and mortality-driven interaction mechanisms.

By identifying a key factor that tunes interaction-driven selection forces, our work informs the design and interpretation of experimental studies. Several recent experimental evolution studies have set out to measure how interspecific interactions affect evolution [6, 7, 8, 9, 10, 40, 41, 42], yet, in the absence of clear expectations for when such effects would be strong, experimental decisions are often arbitrary, and results lack context. Differences between experimental systems might, to some degree, be explained by the growth dynamics in these systems, which range in orders of magnitude (e.g. *D* = 1.1 [8] to *D* = 1500 [43]). However, further work is needed to quantitatively assess this claim across experiments, as the lack of standard reporting currently makes such analysis challenging.

Throughout, we quantify selection once the resident community has reached its ecological steady-state. Away from steady-state, selection can differ: fitness depends not only on which species are present but also on their current abundances, and therefore on the starting composition. This complicates direct comparison with recent efforts to measure how species reshape each other’s distribution of fitness effects [11, 13, 14, 44], which often quantify fitness at a composition set by the experimenter rather than at an ecological equilibrium. Extending our results beyond steady-state will require non-equilibrium approaches; we nonetheless expect the early-bird effect itself to persist, as it reflects a relative robustness to species abundances.

Moreover, considering only steady-state conditions amounts to assuming a separation of timescales between ecology and evolution [45], which entails analyzing only stably coexisting pairs, excluding cases of ‘evolutionary-rescue’ from our study. Yet, experimental works with microbial populations have shown that beneficial mutations can arise quickly and affect population dynamics on a timescale similar to that of ecology [46, 47, 48], further stressing the need to understand these effects outside of steady-state.

Our framework applies most directly when species physiology is stable along a growth cycle—that is, species do not change what they do when conditions change. Many microbial interactions are instead driven by physiological shifts that can be induced by stress-responses, resource concentrations (e.g., diauxic growth), or quorum-sensing [49, 50, 36, 43, 51]. In these cases, species interactions are qualitatively different in a dilution-growth cycle than in a chemostat; a species might secrete a toxin only in stationary phase, which a chemostat never reaches, or two species might compete only once a shared resource is exhausted due to diauxic growth. These kinds of dynamics are not considered in our model. However, our results do provide some intuition for these cases—as long as a growth affecting interaction peaks after the affected species have undergone most of their growth, the interaction will have a small effect relative to early-bird gains.

Our work identifies a key environmental feature — the growth regime — that tunes how strongly species interactions shape evolution, and links this effect to a simple mechanistic cause. The value of the growth regime as a predictor lies in its coarse-grained nature: it applies across diverse ecological systems and is relatively easy to estimate, yet it still yields testable predictions. Specifically, that boom-and-bust systems should exhibit weaker signatures of growth-mediated coadaptations, such as interaction-driven niche differentiation, while retaining or even amplifying mortality-mediated ones, such as the evolution of antitoxins. Other coarse-grained environmental features with comparable broad implications for co-adaptation likely exist, and identifying them offers a promising path toward understanding and predicting short-term evolutionary adaptation.

## Methods

### Model description

#### Resource competition

Two species compete for two substitutable resources. Within each growth cycle, there is no flow (*d* = 0), and dynamics are governed by:

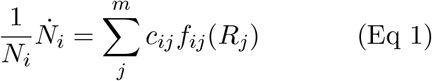

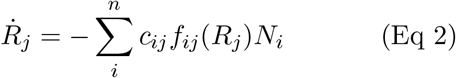

where *N*_*i*_ is the abundance of species *i, R*_*j*_ is the concentration of resource *j, c*_*ij*_ is the maximum per-capita consumption rate of species *i* on resource *j*, and *f*_*ij*_(*R*_*j*_) is the consumption function (linear or Monod, Eq S3, Eq S4). At the end of each cycle, both species and resources are diluted by *D*, with resources replenished with supply concentrations by 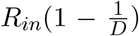. For more details, refer to Section S1.

#### Toxin-mediated interactions

As in resource competition, each growth cycle has no flow (*d* = 0), but here the focal species (species 1) consumes only resources 1 and the secretor (species 2) consumes only resource 2. The secretor produces a toxin *T* during growth at a rate proportional to its consumption:

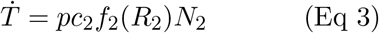

where *p* is the per-unit-growth toxin production coefficient. The toxin affects the focal species through one of two mechanisms:

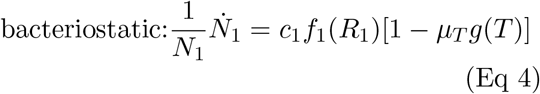

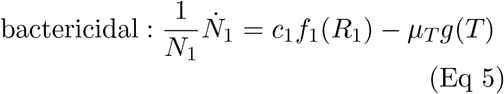

where *µ*_*T*_ is the maximum toxin effect (growth reduction or kill rate, respectively) and *g*(*T*) is a Hill function:

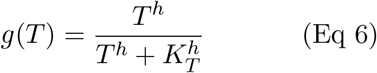

Resource and secretor dynamics follow the same form as described for resource competition. Dilution at cycle ends is applied to species, resources, and toxin alike. For more details, refer to Section S6.

### Simulating growth dynamics

We simulated growth dynamics under dilution-growth conditions as described in Section S2, for both resource competition and toxin-mediated interference. In all simulations, each growth cycle was integrated independently using an explicit Runge-Kutta method (`scipy.integrate.solve_ivp` with RK45) in Python, and dilution by factor *D* was applied between cycles.

Each simulation was run for a duration sufficient for the system to reach steady state, defined as identical dynamics across consecutive cycles (simulation lengths are reported in Fig. S3, and Fig. S5). We excluded simulations in which (i) one of the species was excluded, defined as reaching an abundance below 10^*−*4^, or (ii) the system did not reach steady state, defined as | *v*^*f*^ (0) *™ v*^*f™*1^(0) |*>* 10^*−*8^, where *v*(0) denotes the abundance of any species or resource at the beginning of the cycle and *f* is the last cycle that was simulated.

#### Resource competition

For each randomly drawn parameter set, we simulated the dynamics across 10 values of *τ* (listed in S1), adjusting *D* accordingly to hold the time-averaged dilution rate *d* constant (see S2.2).

#### Toxin-mediated interactions

We drew 300 parameter sets and simulated each under three conditions: focal monoculture, coculture with a bacteriostatic toxin, and coculture with a bactericidal toxin, all at *τ* = 100. From these, we randomly selected 30 parameter sets and re-simulated each across 4 values of *τ* (listed in S3) for the analysis shown in Figures 5B and 5C.

### Introducing mutations

Once a simulation reached steady state, we introduced mutants into the system to measure invasion fitness. For each simulation, we introduced 30 independent mutations, each tested separately in both the monoculture and the coculture. Mutants were introduced at an abundance of 10^*−*5^, and inherited the focal species’ parameters with small random perturbations, such that:

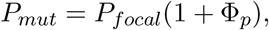

where *p* is a parameter and Φ_*p*_ is drawn independently for each mutated parameter from a uniform distribution between *−*0.1 and 0.1.

Unless stated otherwise, parameters were mutated jointly: *c*_11_, *c*_12_, *k*_11_, *k*_12_ for resource competition, with the additional max toxin effect *µ*_*T*_ included for toxin simulations. The exception is the analysis in Figures 5B and 5C, where we used a specific tradeoff mutation with a fixed +20% change in maximum growth rate (*c*_11_) and a fixed +50% change in maximum toxin effect (*µ*_*T*_, see S5 for more detail).

We measured invasion fitness over a single growth cycle as the difference between the mutant’s and the focal ancestor’s per-capita growth rates:

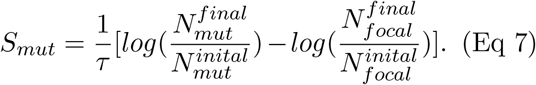

### Quantifying interactions

In order to quantify the temporal dynamics of interactions we use the *instantaneous interaction* metric developed in [36]. For a pair of species interacting solely through resource competition, it is simplified to:

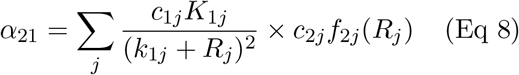

*α*_21_ quantifies the per-capita effect of species 2 on species 1. This could then be transformed into the total instantaneous interaction by multiplying by the population size of species 2, *α*_21_*N*_2_, and to the cumulative interaction by integrating *α*_21_*N*_2_ over the course of a cycle. To quantify interactions over the course of a cycle, we used simulation data with a 0.5 timestep spacing and integrated using `scipy.cumulative_trapezoid`.

## Acknowledgments

We thank Amir Erez for providing valuable feedback on the manuscript. We thank the Friedman lab and the Microbial Systems Ecology group for insightful discussions. This work was supported by the Hebrew University Center for Sustainability and the Israeli Science Foundation (grant number 883/22).

## Notes

### Competing Interest Statement

The authors have declared no competing interest.

