## Supplementary Information for "Early bird dynamics reduce selection imposed by species interactions in boom-and-bust regimes"

**S7 Simulating toxin-mediated interactions** 19

#### S1 Consumer-resource model

We consider a community of $n$ species competing for $m$ substitutable resources. The dynamics of species $i$ and resource $j$ are given by:

$$\frac{1}{N_i}\dot{N}_i = \sum_{j=1}^{m} c_{ij} f_{ij}(R_j) - d(t), \tag{Eq S1}$$

$$\dot{R}_j = d(t)(R_{j,\mathrm{in}} - R_j) - \sum_{i=1}^{n} c_{ij} f_{ij}(R_j) N_i, \tag{Eq S2}$$

where $N_i$ is the abundance of species $i$, $R_j$ is the concentration of resource $j$, $c_{ij}$ is the maximal consumption rate of species $i$ on resource $j$, $f_{ij}(R_j)$ is the consumption function describing the dependence of consumption on resource concentration, $d(t)$ is the flow rate, and $R_{j,\mathrm{in}}$ is the supply concentration of resource $j$. The per-capita growth rate of species $i$ is the sum of its consumption rates across all available resources. Each resource is consumed independently, and the consumed resources are removed from the environment and converted to the biomass of the consuming species.

**Consumption functions.** We consider two forms for the consumption function $f_{ij}(R_j)$:

1. **Linear:**

$$f_{ij}(R_j) = R_j. \tag{Eq S3}$$

   In this case, consumption increases linearly with resource concentration without saturation.

2. **Monod (saturating):**

$$f_{ij}(R_j) = \frac{R_j}{k_{ij} + R_j}, \tag{Eq S4}$$

   where $k_{ij}$ is the half-saturation constant of species $i$ for resource $j$. Consumption saturates at high resource concentrations, approaching the maximal rate $c_{ij}$.

#### S2 Growth regimes

We model population and resource dynamics as a series of dilution-growth cycles, and use this framework to describe a continuum of growth regimes; from continuous, chemostat-like flow to pronounced intermittent flow that leads to boom-and-bust dynamics. In this framework, populations and resources experience a growth phase of duration $\tau$ during which there is no in- or out-flow ($d(t) = 0$) and dynamics are governed by consumption and growth alone, followed by an instantaneous dilution event at the end of each cycle in which populations are diluted by a factor $D$ and resources are replenished from the supply at concentration $R_{j,in}$. The species and resource concentrations at the start of cycle $n+1$ are therefore related to those at the end of cycle $n$ by:

$$N_i^{n+1}(0) = \frac{1}{D} N_i^n(\tau), \tag{Eq S5}$$

$$R_j^{n+1}(0) = \frac{1}{D} R_j^n(\tau) + \frac{(D-1)}{D} R_{j,in}. \tag{Eq S6}$$

In this setup, a steady state is defined as a state in which resource and population dynamics within each cycle are identical; therefore:

$$N_i^n(0) = N^{n+1}(0) = \frac{1}{D} N_i^n(\tau), \tag{Eq S7}$$

$$R_j^n(0) = R^{n+1}(0) = \frac{1}{D} R_j^n(\tau) + \frac{(D-1)}{D} R_{in}. \tag{Eq S8}$$

Dropping the cycle index, we get that in every cycle:

$$N_i(\tau) = D N_i(0), \tag{Eq S9}$$

$$R_j(0) = \frac{R_j(\tau) + (D-1) R_{j,in}}{D}. \tag{Eq S10}$$

Another quantity of interest is the average per-capita growth over a cycle. The per-capita growth of species $i$ is denoted as $\gamma_i(\mathbf{R}) = \frac{1}{N_i}\dot{N}_i$, where $\mathbf{R} = (R_1, R_2, \ldots, R_m)$, and its average over a cycle is given by:

$$\langle \frac{1}{N_i} \frac{dN_i}{dt} \rangle = \langle \gamma_i(\mathbf{R}) \rangle = \frac{1}{\tau} \int_0^\tau \gamma_i(\mathbf{R}) dt. \tag{Eq S11}$$

Using the fact that

$$N_i(\tau) = N_i(0) e^{\int_0^\tau \gamma_i(\mathbf{R})} dt,$$

we get that at steady state:

$$\langle \gamma_i(\mathbf{R}) \rangle = \frac{1}{\tau} \int_0^\tau \gamma_i(\mathbf{R}) dt = \frac{1}{\tau} \log \frac{N_i(\tau)}{N_i(0)} = \frac{1}{\tau} \log D. \tag{Eq S12}$$

##### S2.1 The chemostat limit

These two regimes could be studied as a continuum if we describe the chemostat as the limit where the time between dilutions ($\tau$) tends to 0 and the dilution factor ($D$) tends to 1. However, care is needed in taking this limit to ensure that the average nutrient supply and dilution rates do not change. For simplicity, in the following derivations, we will focus on a single species and a single nutrient, and drop the corresponding subscript. Selection in the case of growth on multiple resources is considered in Section S3.1.

Specifically, we need to ensure that, at steady state, the average dilution rate in the chemostat $\bar{d}$ is equal to the average per-capita growth rate over a growth cycle. Therefore, the relationship between the species and resource concentrations at the end of a cycle and at the beginning of the next cycle is given by:

$$\overline{d} = \langle cf(R) \rangle = \frac{1}{\tau} \log D. \quad \text{(Eq S13)}$$

Taking the limit $\tau \to 0$ while adjusting $D$ according to this relationship recovers the chemostat model with flow rate $\overline{d}$.

Note that while the average per capita growth rate of any species at steady state will be independent of the batch parameters $(\tau, D)$, that is generally not the case for other quantities. For example, when consumption is non-linear, or when multiple resources are consumed simultaneously (even linearly), the average resource amount varies with the batch parameters even when $d$ is held constant.

#### S2.2 The boom-and-bust limit

The opposite situation to the chemostat limit is when each batch is long and is followed by a very large dilution. In this case, the initial resource abundance at each cycle tends to the input concentration

$$R(0) = \frac{R(\tau) + (D-1)R_{in}}{D} \overset{D \gg 1}{\approx} R_{in},$$

and approximately all the resources are consumed and converted to biomass during the cycle. Therefore:

$$R(\tau) \approx 0 \quad \text{(Eq S14)}$$

$$N(\tau) \approx N(0) + R_{in} \approx R_{in}. \quad \text{(Eq S15)}$$

Since the initial species' abundance is small, it's depletion of the resource is initially negligible, and its growth is exponential at an approximately constant rate until the resources are significantly depleted. Due to the species' exponential growth, most of the resource consumption occurs over a short time interval compared to the length of a cycle. Therefore, we can approximate the resource dynamics as a step function, going from $R_{in}$ to 0 at some time $\hat{t}$. That is, we'll use the following approximation:

$$N(t) \approx \begin{cases} N(0)e^{\gamma(R_{in})t} & t \leq \hat{t} \\ N_\tau & t > \hat{t}, \end{cases} \quad \text{(Eq S16)}$$

$$R(t) \approx \begin{cases} R_{in} & t \leq \hat{t} \\ 0 & t > \hat{t}. \end{cases} \quad \text{(Eq S17)}$$

The depletion time $\hat{t}$ can be found by solving $N(\tau) = N(\hat{t})$:

$$\hat{t} = \frac{1}{\gamma(R_{in})} \log \frac{N(\tau)}{N(0)} = \frac{1}{\gamma(R_{in})} \log D. \quad \text{(Eq S18)}$$

Under this approximation, the temporal average of any function of the resource $(g(R))$ can be readily computed:

$$\langle g(R) \rangle = \frac{1}{\tau} \int_0^\tau g(R)\, dt \approx \frac{1}{\tau} \int_0^{\hat{t}} g(R)\, dt \approx \frac{1}{\tau} g(R_{in})\hat{t} = \frac{g(R_{in})}{\gamma(R_{in})} \frac{1}{\tau} \log D = \frac{g(R_{in})}{\gamma(R_{in})} \overline{d}. \quad \text{(Eq S19)}$$

#### S3 Invasion fitness in batch culture

##### S3.1 General derivation

We define a mutant's invasion fitness over a single growth cycle as the difference between the mutant's and its ancestor's per-capita growth rates:

$$s = \langle \gamma_{\text{mut}}(\mathbf{R}) \rangle - \langle \gamma(\mathbf{R}) \rangle. \tag{Eq S20}$$

Plugging in Eq S12, we get that the invasion fitness can also be expressed as the difference between the time-averaged number of generations over a cycle:

$$s = \frac{1}{\tau} \left[ \ln \frac{N_{\text{mut}}(\tau)}{N_{\text{mut}}(0)} - \ln \frac{N(\tau)}{N(0)} \right]. \tag{Eq S21}$$

In some cases, invasion fitness can be independent of the batch parameters and identical to that of the equivalent chemostat case (i.e. dependent only on $\overline{d}$, and not on $\tau$ or $D$).

One such notable case is when the mutant per capita growth rate is different from the ancestor's by a fixed amount $\Delta$ that is independent of resource concentrations:

$$\gamma_{\text{mut}}(\mathbf{R}) = \gamma(\mathbf{R}) + \Delta. \tag{Eq S22}$$

The mutant's invasion fitness is then simply:

$$s = \langle \Delta \rangle = \Delta. \tag{Eq S23}$$

Another notable case is when the mutant per capita growth rate is different from the ancestor's by a fixed proportion:

$$\gamma_{\text{mut}}(\mathbf{R}) = (1 + \Delta)\gamma(\mathbf{R}). \tag{Eq S24}$$

In this case, the mutant's invasion fitness is again independent of the batch parameters and is given by:

$$s = \langle \Delta\gamma(\mathbf{R}) \rangle = \Delta \langle \gamma(\mathbf{R}) \rangle = \Delta \overline{d}. \tag{Eq S25}$$

More generally, if the mutant's growth rate is a linear function of the ancestor's growth rate, the invasion fitness will only be a function of the ancestor's average growth rate ($\overline{d}$), which is independent of the batch parameters. However, invasion fitness will depend on batch parameters if that is not the case. For example, invasion fitness may differ from the chemostat limit if there are multiple resources, even if they are all consumed linearly (see below).

##### S3.2 Invasion fitness in the chemostat limit

At the chemostat limit, the resource level is approximately fixed at the steady state level of the resident's steady state $\mathbf{R}^*$. Therefore, the invasion fitness in the chemostat limits is given by:

$$s_{D \simeq 1} = \langle \gamma_{\text{mut}}(\mathbf{R}) \rangle - \langle \gamma(\mathbf{R}) \rangle = \gamma_{\text{mut}}(R^*) - \gamma(R^*), \tag{Eq S26}$$

and fitness depends on growth rates at the resident's steady-state resource levels.

##### S3.3 Invasion fitness in the boom-and-bust limit

At the boom-and bust limit (large $D$ limit), for a single resource ($R$), using [Eq S19] the invasion fitness is given by:

$$s_{D \gg 1} = \langle \gamma_{\text{mut}}(R) \rangle - \overline{d} \approx \frac{\gamma_{\text{mut}}(R_{in})}{\gamma(R_{in})} \overline{d} - \overline{d} = \left( \frac{\gamma_{\text{mut}}(R_{in}) - \gamma(R_{in})}{\gamma(R_{in})} \right) \overline{d} \tag{Eq S27}$$

$$= \frac{\overline{d}}{\gamma(R_{in})} \left( \gamma_{\text{mut}}(R_{in}) - \gamma(R_{in}) \right). \tag{Eq S28}$$

When there are multiple resources (substitutable and simultaneously consumed), if the mutant's overall per capita growth rate changes in an absolute or proportional manner, selection is dependent only on the average dilution rate, as in the single resource case (see Section S3.1). However, when mutations affect on the consumption rate of each resource differently, selection in general does depend on the specific batch duration and dilution factor.

As in the single resource case, we'll use the approximation that all resources are at their input level ($\mathbf{R}_{in}$ ) until a time ($\hat{t}$ ) when they are all rapidly consumed.

Note that it may be possible to derive a more accurate piece-wise approximation where resources are depleted sequentially and the exponential growth rate decreases as each resource is depleted. However, this approximation is often quite accurate Fig.??B, as depletion times are typically very close, as they are driven by the the increase in abundance of the shared consumer.

Under this approximation the invasion fitness is directly analogous to the single resource case [??]

$$s_{D\gg 1} = \left(\frac{\gamma_{\text{mut}}(\mathbf{R_{in}}) - \gamma(\mathbf{R_{in}})}{\gamma(\mathbf{R_{in}})}\right)\overline{d}. \qquad \text{(Eq S29)}$$

**Mutations that affect growth on specific resources**

We'll now consider some specific cases in which mutation have an additive or proportional effect on the growth rate that is independent of resource concentrations, but these effects can differ between resources. In the additive case, even if the mutant has a different growth effect on each resource, it's invasion fitness is independent of the batch parameters:

$$\gamma_{\text{mut}}(\mathbf{R}) = \sum_i^L f_{\text{mut,i}}(R_i) = \sum_i^L (f_{\text{i}}(R_i) + \Delta) = \gamma(\mathbf{R}) + \Delta, \qquad \text{(Eq S30)}$$

$$s = \langle\gamma(\mathbf{R})\rangle + \Delta - \overline{d} = \langle\Delta\rangle = \Delta. \qquad \text{(Eq S31)}$$

In contrast, in the case of proportional effects, the invasion fitness does depend on average consumption rates over a batch:

$$\gamma_{\text{mut}}(\mathbf{R}) = \sum_i^L f_{\text{i}}(R_i)(1 + \Delta_i) = \gamma(\mathbf{R}) + \sum_i^L f_{\text{i}}(R_i)\Delta_i, \qquad \text{(Eq S32)}$$

$$s = \sum_i^L \langle f_{\text{i}}(R_i)\rangle\Delta_i. \qquad \text{(Eq S33)}$$

When each resource is consumed linearly ($f_i(R_i) = c_iR_i$), one may think that $s$ would be independent of the batch parameters since it depends only on the batch-average resource amount. However, even in this simple case the average resource levels do vary with batch parameters and are dependent on each other since they are coupled through the population size:

$$\langle c_iR_i\rangle = \frac{1}{\tau}\int_0^\tau c_iR_i\,dt = \frac{1}{\tau}\int_0^\tau \frac{c_iR_i}{N\sum_j c_jR_j}\,dN. \qquad \text{(Eq S34)}$$

Therefore, even in this relatively simple case, it not not possible to derive a general analytical expression for the invasion fitness. However, at the boom-and-bust limit, the invasion fitness in this case can be approximated as:

$$s_{D\gg 1} = \frac{\sum_i^L f_i(R_{in,i})\Delta_i}{\gamma(\mathbf{R_{in}})}\overline{d} = \frac{\sum_i^L f_i(R_{in,i})\Delta_i}{\sum_j^L f_j(R_{in,j})}\overline{d}. \qquad \text{(Eq S35)}$$

#### S4 Simulating selection under resource competition

In order to study the differences in how resource competition affects selection in chemostat vs boom-and-bust regimes, we simulate the invasion of random mutants into steady-state monoculture and co-cultures.

For this, we conduct the following workflow:

1. Choose random parameters and simulate in chemostat. Random parameters include those of the species (max growth rate $c$, half saturation constant $k$), and environmental parameters (flow in resource concentrations $R_{in}$, flow rate $\overline{d}$). see S1 for the distributions from which these are taken.

2. Run batch simulation for each parameter combination across $\tau$ values, keeping $d$ constant using Eq S13. Batch simulations are only conducted on pairs that coexist, defined by both species keeping an abundance above $10^{-4}$. In case the steady-state could be computed analytically, this step is skipped.

3. Seed a random mutants and simulate for one cycle. Compute the selection using Eq S21

##### S4.1 One linearly consumed resource

As already well established[1], fluctuations in death rate and resource availability do not affect selection if the growth rate is determined by one resource whose consumption function is linear. Our simulations indeed follow this rule (Fig S1).

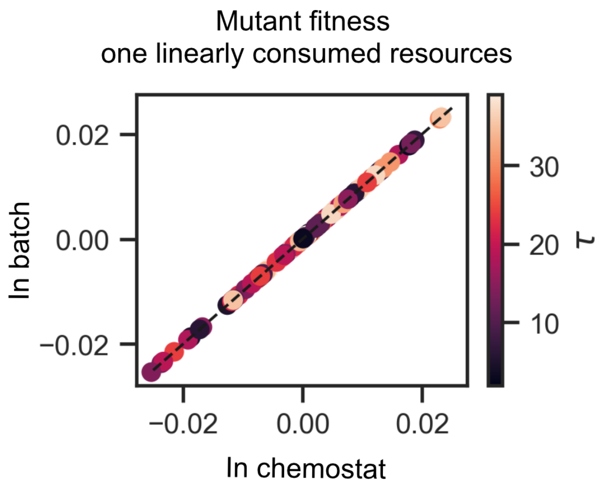

**Figure S1: Selection is not affected by growth regime on one linearly consumed resource**. The invasion fitness of mutants in a dilution-growth regime with varying cycle lengths ($\tau$, color scheme), against the fitness in the corresponding chemostat. The plot shows the results of 100 simulations with randomly drawn max growth rate ($c_1 \sim \mathcal{U}(2, 4)$), flow rate ($\overline{d} \sim \mathcal{U}(0.01, 3)$), and cycle length ($\overline{d} \sim \mathcal{U}(2, 40)$).

##### S4.2 Two linearly consumed resources

The equivalence between the chemostat and the boom-and-bust regime breaks as soon as there is more than one resource, even if the resources are consumed linearly (S2A). This results in an increased similarity in selection between monoculture and coculture in the boom-and-bust regime (S2B).

##### S4.3 Parameter distributions and pipeline

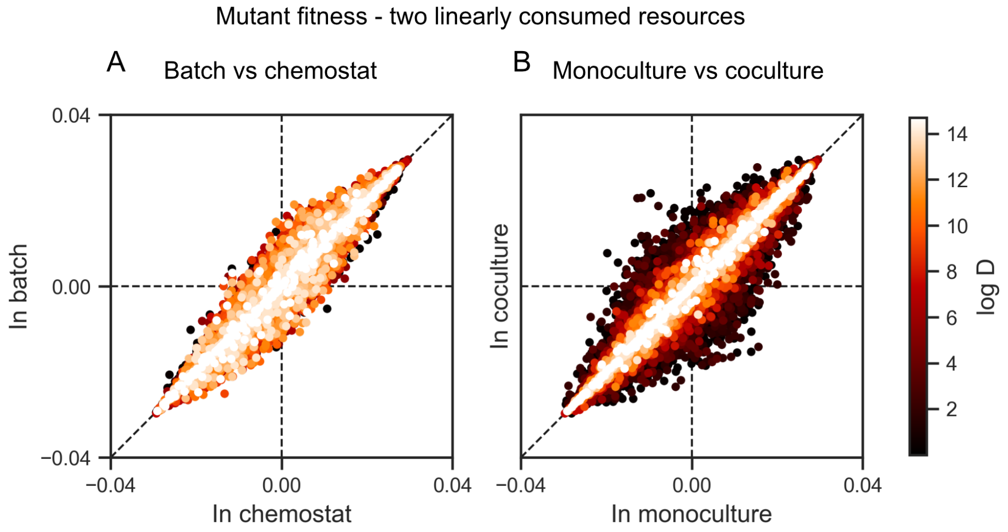

**Figure S2: Selection differs between chemostat and boom-and-bust cycles on linearly consumed resources**. Mutant invasion fitness at steady state in (A) batch dynamics vs the equivalent chemostat, (B) monoculture vs coculture. Points represent 30 mutations across 1913 unique simulations, each evaluated under 9 $\tau$ values. Point color denotes the strength of the boom-and-bust dynamics, mapped to the log dilution factor $D$.

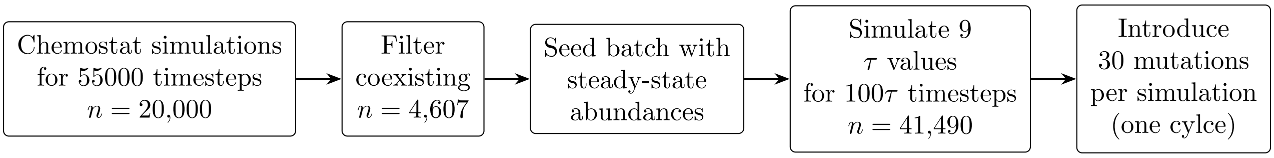

**Figure S3:** Simulation pipeline for resource competition.

Table S1 : Parameter distributions and simulation counts for resource competition.

| **Parameter** | **Distribution / Value** | **Notes** |
|---|---|---|
| *Species parameters (per simulation, drawn independently)* | | |
| $C_{ij}$ (consumption rate) | Uniform$[0.1, 5.0]$ | 4 values per simulation |
| $k_{ij}$ (half-saturation) | Log-uniform on $\{10^6, \ldots, 10^9\}$, | 10 values |
| *Environmental parameters* | | |
| $R_{j,\text{in}}$ (supply) | Log-spaced, $10^6$ to $10^9$ (100 values) | Drawn independently per resource |
| $\bar{d}$ (avg. dilution rate) | Linear, $[0.001, 0.3]$ (20 values) | Held constant across $\tau$ |
| *Growth regime* | | |
| $\tau$ (cycle duration) | $\{1, 7, 13, 19, 25, 31, 37, 43, 49\}$ | 9 values |
| $D$ (dilution factor) | $\exp(\bar{d}\tau)$ | Set to hold $d$ constant |
| *Simulation counts* | | |
| Chemostat seed simulations | 20,000 | 1000 per $d$ value |
| Coexisting at steady state | 4,610 | Used to seed batch |
| Batch simulations | 41,490 | 4,610 $\times$ 9 $\tau$ values |
| *Mutations* | | |
| Number of mutations | 30 | |
| Mutated parameters | $C_{ij}$, $k_{ij}$ | |
| Distribution | uniform[-0.1, 0.1] | |

### S5 The evolutionary early-bird effect

Here we show why mutations affecting growth at higher resource concentrations result in stronger selection than equivalent mutations acting at lower concentrations. We first consider one resource and then extend the result to simultaneous consumption of multiple substitutable resources. As in other sections, we assume the mutant is rare enough that it does not affect the population or resource trajectory.

#### S5.1 Growth on a single resource

During the growth phase, let the ancestor's per-capita growth rate be $g(R) = cf(R)$. The within-batch dynamics simplify to

$$\dot{N} = Ng(R), \tag{Eq S36}$$

$$\dot{R} = -Ng(R). \tag{Eq S37}$$

Consider a mutant with growth rate

$$g_{\mathrm{mut}}(R) = [1 + \epsilon(R)]g(R), \tag{Eq S38}$$

where $\epsilon(R)$ is the fractional change in growth rate caused by the mutation at resource concentration $R$. Its invasion fitness over one cycle is

$$S_{\mathrm{mut}} = \frac{1}{\tau}\int_0^\tau [g_{\mathrm{mut}}(R) - g(R)]\, dt = \frac{1}{\tau}\int_0^\tau \epsilon(R)g(R)\, dt. \tag{Eq S39}$$

Using $-dR = Ng(R)dt$ gives

$$S_{\mathrm{mut}} = \int_{R(\tau)}^{R(0)} \epsilon(R)w(R)\, dR, \qquad w(R) = \frac{1}{\tau N(R)}. \tag{Eq S40}$$

Thus, $w(R)$ is the contribution to selection per unit resource concentration for a fixed fractional effect $\epsilon(R)$. Since

$$\frac{dN}{dR} = -1, \tag{Eq S41}$$

we have

$$\frac{dw}{dR} = \frac{1}{\tau N^2} > 0. \tag{Eq S42}$$

The same change in growth over the same concentration range therefore contributes more to selection when it occurs at higher $R$. At periodic steady state, $N(\tau) = DN(0)$, so the weight at the beginning of the cycle is $D$ times that at the end.

**Equal effects over different concentration intervals.** To illustrate the larger contribution to the selection of a mutation affecting growth at higher resource concentrations, consider a mutation that changes growth by the same fraction $\epsilon_0$ over a concentration interval of width $\Delta R$, whose lower edge is $R_L$:

$$\epsilon(R) = \begin{cases} \epsilon_0, & R_L \le R \le R_L + \Delta R, \\ 0, & \text{otherwise}. \end{cases} \tag{Eq S43}$$

Its invasion fitness is

$$S_{\mathrm{mut}}(R_L) = \frac{\epsilon_0}{\tau}\int_{R_L}^{R_L+\Delta R} \frac{dR}{N(R)}. \tag{Eq S44}$$

For mutations with the same $\epsilon_0$ and $\Delta R$,

$$\frac{\partial S_{\mathrm{mut}}}{\partial R_L} = \frac{\epsilon_0}{\tau}\left[\frac{1}{N(R_L + \Delta R)} - \frac{1}{N(R_L)}\right]. \tag{Eq S45}$$

Because $N(R)$ decreases with $R$, this derivative has the sign of $\epsilon_0$. Thus, for beneficial mutations ($\epsilon_0 > 0$), moving the affected interval to higher resource concentrations increases selection; for deleterious mutations, it increases the magnitude of negative selection.

**Monod affinity mutation.** The early-bird weighting does not imply that the fractional effect of a mutation is itself largest at high resource concentration. Consider an ancestor and a mutant with the same maximum growth rate $c$ but different Monod half-saturation constants,

$$g_F(R) = c\frac{R}{k_F + R}, \qquad \text{(Eq S46)}$$

$$g_M(R) = c\frac{R}{k_M + R}. \qquad \text{(Eq S47)}$$

The fractional growth-rate difference is

$$\epsilon(R) = \frac{g_M(R) - g_F(R)}{g_F(R)} = \frac{k_F - K_M}{k_M + R}. \qquad \text{(Eq S48)}$$

For an affinity-improving mutation, $k_M < k_F$, $\epsilon(R)$ increases as $R$ decreases. Selection over a dilution-growth cycle nevertheless follows Eq. Eq S40, and therefore combines this stronger effect at low $R$ with the larger evolutionary weight of high $R$.

#### S5.2 Multiple substitutable resources

Now consider a focal population consuming $m$ substitutable resources, with

$$g_F(\mathbf{R}) = \sum_{j=1}^{m} c_j f_j(R_j), \qquad \text{(Eq S49)}$$

and

$$\dot{R}_j = -N c_j f_j(R_j). \qquad \text{(Eq S50)}$$

Suppose a mutation affects only growth supported by resource $R_k$:

$$g_M(\mathbf{R}) = g_F(\mathbf{R}) + \epsilon_k(R_k) c_k f_k(R_k), \qquad \text{(Eq S51)}$$

where $\epsilon_k(R_k)$ is the fractional change in the growth contribution from resource $k$. Its invasion fitness is

$$S_{\text{mut}} = \frac{1}{\tau}\int_0^{\tau} \epsilon_k(R_k) c_k f_k(R_k)\, dt = \int_{R_k(\tau)}^{R_k(0)} \epsilon_k(R_k) w_k(R_k)\, dR_k, \qquad \text{(Eq S52)}$$

with

$$w_k(R_k) = \frac{1}{\tau N(R_k)}. \qquad \text{(Eq S53)}$$

While $R_k$ is being consumed,

$$\frac{dN}{dR_k} = -\frac{g_F(\mathbf{R})}{c_k f_k(R_k)} < 0, \qquad \text{(Eq S54)}$$

and therefore

$$\frac{dw_k}{dR_k} = \frac{g_F(\mathbf{R})}{\tau N^2 c_k f_k(R_k)} > 0. \qquad \text{(Eq S55)}$$

Thus, simultaneous consumption of other resources does not remove the early-bird weighting for a mutation affecting one resource: equal concentration intervals of $R_k$ receive greater weight at higher $R_k$.

#### S5.3 Assumptions and limits

The derivations above use the unit-yield convention of Eq S2. Different resources may have different constant yields without changing the result. If resource $j$ has yield $Y_j$, then $\dot{R}_j = -Nc_j f_j(R_j)/Y_j$ and the weight becomes $w_j(R_j) = Y_j/[\tau N(R_j)]$. The constant $Y_j$ can equivalently be absorbed into the units of $R_j$, and does not affect the increase of $w_j$ with resource concentration.

We have considered only mutations affecting the growth rate on a single resource at a time. If a mutation affects several resources simultaneously, each contribution can be written separately in the form above, but the net selection depends on their relative effects and on the joint resource trajectory. Diauxic or regulated resource use can also decouple high resource concentration from the beginning of the cycle: a resource may remain abundant until the population is already large. The monotonic result still applies while that resource is actively consumed, but mutations affecting switching or regulation are not described by the derivation above.

Finally, if other species also consume the focal resource, its depletion is no longer determined by the focal population alone. For resource $R_k$,

$$-\dot{R}_k = \sum_i N_i c_{ik} f_{ik}(R_k), \qquad \text{(Eq S56)}$$

so a mutation affecting the focal species' use of $R_k$ has weight

$$w_k(R_k) = \frac{c_{Fk} f_{Fk}(R_k)}{\tau \sum_i N_i c_{ik} f_{ik}(R_k)}. \qquad \text{(Eq S57)}$$

This weight need not increase monotonically with $R_k$. A competitor can affect resource concentrations at which growth advantages contribute most to selection in a focal species.

### S6 Toxin-mediated interactions

To model interactions that are not mediated by shared resources, we consider a scenario in which two species each consume a separate resource but one species (the secretor, species 2, consumes resource 2) produces a toxin $T$ that affects the other (the focal species, species 1, consumes resource 1).

#### S6.1 Bactericidal toxin

First, we consider a bactericidal toxin, that kills the focal species. The within-batch dynamics are given by:

$$\frac{1}{N_1}\dot{N}_1 = c_1 f_1(R_1) - \mu g(T) \qquad \text{(Eq S58)}$$

$$\frac{1}{N_2}\dot{N}_2 = c_2 f_2(R_2) \qquad \text{(Eq S59)}$$

$$\dot{R}_i = -c_i f_i(R_i) N_i \qquad \text{(Eq S60)}$$

$$\dot{T} = pP(N_2). \qquad \text{(Eq S61)}$$

We focus on selection on species 1 that is being killed by the toxin ($T$) produced by species 2. For simplicity, we'll consider only mutations that affect the maximal growth rate ($c_1$) and the maximal death rate ($\mu$). Selection for such mutants is given by:

$$s = \langle \gamma_m - \gamma \rangle = (c_m - c_1)\langle f_1(R_1)\rangle - (\mu_m - \mu)\langle g(T)\rangle \qquad \text{(Eq S62)}$$

**Chemostat limit**

In order to find an expression fo the selection in the chemostat, we'll first find the steady-state levels of the resource consumed by the focal species ($R_1$) and the toxin ($T$).

In monoculture:

$$T^{(m)} = 0$$
$$f_1(R_1^{(m)}) = \frac{\overline{d}}{c_1},$$

where $^{(m)}$ denotes the steady-state of species 1 in monoculture.

In coculture there is both more resource and more toxin at steady-state:

$$T^{(c)} = \frac{pP(N_2^{(c)})}{d} \qquad \text{(Eq S63)}$$

$$f_1(R_1^{(c)}) = \frac{\overline{d}}{c_1} + \frac{\mu}{c_1} g(T^{(c)}) = f_1(R_1^{(m)}) + \frac{\mu}{c_1} g(T^{(c)}). \qquad \text{(Eq S64)}$$

Using these steady-state value, we get that selection in mono- and in coculture is given by:

$$S_{D\simeq 1}^{(m)} = \frac{c_m - c_1}{c_1}\overline{d} \qquad \text{(Eq S65)}$$

$$S_{D\simeq 1}^{(c)} = (c_m - c_1)\left(\frac{\overline{d}}{c_1} + \frac{\mu g(T^{(c)})}{c_1}\right) - (\mu_m - \mu) g(T^{(c)}). \qquad \text{(Eq S66)}$$

We can express the selection in coculture as:

$$S_{D\simeq 1}^{(c)} = S_{D\simeq 1}^{(m)} + g(T^{(c)})\left(\frac{c_m}{c_1}\mu - \mu_m\right). \qquad \text{(Eq S67)}$$

For mutations that only change $c_1$ this simplifies to:

$$S^{(c)}_{D\simeq 1} = S^{(m)}_{D\simeq 1} + g(T^{(c)})\frac{c_m - c_1}{c_1}\mu. \quad \text{(Eq S68)}$$

Note that selection is different from monoculture even though the mutation doesn't affect antibiotic sensitivity. This is because the steady-state resource levels in coculture are higher than in monoculture due to the toxin reducing the steady-state levels of the focal species, elevating the concentration of the resource this population consumes.

**Boom-and-bust limit ($D \gg 1$)**

In monoculture, selection is independent of the batch parameters and is identical to the chemostat limit. We've derived this result in previous sections for the maximal growth rate ($c_1$), and the maximal mortality rate ($\mu$) is trivially not under selection in any monoculture conditions since no toxin is present in the system. That is, for mutations affecting these traits

$$s^{(m)}_{D\gg 1} = s^{(m)}_{D\simeq 1}. \quad \text{(Eq S69)}$$

To compute the selection in batch condition in coculture ($s^{(c)}_{D\gg 1}$), we'll again approximate the growth and resource consumption of each species as a step function. Since each species consumes a different resource, they'll also grow and exhaust their respective resources at different times ($\hat{t}_i$). In addition, the toxin will initially be approximately 0, and then instantaneously increase to a level $T_f$ when the producing species grows (at $\hat{t}_2$).

Selection depends on the averages of the toxin mortality and the resource consumption over a batch. Average toxin mortality is given by:

$$\langle g(T)\rangle \approx \frac{1}{\tau}\int_{\hat{t}_2}^{\tau} g(T)\,dt \approx \frac{1}{\tau}(\tau - \hat{t}_2)g(T_f) = \left(\frac{\tau - \hat{t}_2}{\tau}\right) g(T_f). \quad \text{(Eq S70)}$$

That is, the average toxin mortality is approximately the fraction of the batch length during which the toxin is present times the level of toxin mortality at the final toxin level.

As in the previous section on resource competition, the growth time of species 2 is given by:

$$\hat{t}_2 = \frac{1}{c_2 f(R_{2,\mathrm{in}})}\log D. \quad \text{(Eq S71)}$$

Plugging this into Eq S70 gives

$$\langle g(T)\rangle \approx (1 - \frac{\overline{d}}{c_2 f_2(R_{2,\mathrm{in}})})g(T_f) = \left(1 - \frac{f_2(R_2^{(c)})}{f_2(R_{2,\mathrm{in}})}\right) g(T_f). \quad \text{(Eq S72)}$$

Next, we'll find an expression for the average resource consumption over a batch. To do so, we first need to derive an expression for the growth time of species 1. Under the boom-and-bust approximation, species 1 grows at a constant rate $c_1 f_1(R_{1,\mathrm{in}})$ for a time $\hat{t}_1$ and dies at a constant rate $\mu g(T_f)$ for a time $\tau - \hat{t}_2$. Therefore, the overall growth of species 1 over a batch is given by

$$N_1(\tau) = N_1(0)e^{c_1 f_1(R_{1,\mathrm{in}})\hat{t}_1 - \mu g(T_f)(\tau - \hat{t}_2)}. \quad \text{(Eq S73)}$$

Note that since growth is exponential and these are per-captia growth rates, the order of events does not change the expression. That is, it does not matter whether the toxin is produces before or after species 1 grows.

As before, we can now get the growth time $\hat{t}_1$ by imposing that the species grows by a factor $D$ over the entire batch:

$$\log D = \log\frac{N_1(\tau)}{N_1(0)} = c_1 f_1(R_{1,\mathrm{in}})\hat{t}_1 - \mu g(T_f)(\tau - \hat{t}_2), \quad \text{(Eq S74)}$$

giving

$$\hat{t}_1 = \frac{1}{c_1 f_1(R_{1,\mathrm{in}})}\left[\log D + \mu g(T_f)(\tau - \hat{t}_2)\right]. \tag{Eq S75}$$

The growth times gives the approximate average resource consumption over a batch

$$\langle f_1(R_1)\rangle \approx \frac{\hat{t}_1}{\tau} f_1(R_{1,\mathrm{in}}) \approx \frac{\overline{d}}{c_1} + \frac{ug(T_f)}{c_1}\left(\frac{\tau - \hat{t}_2}{\tau}\right) = f_1(R_1^{(1)}) + \frac{ug(T_f)}{c_1}\left(\frac{\tau - \hat{t}_2}{\tau}\right). \tag{Eq S76}$$

Using these averages, we get an expression for the selection in coculture

$$s^{(c)}_{D\gg 1} \approx s^{(m)} + g(T_f)\left(\frac{c_m}{c_1}\mu - \mu_m\right)\left(\frac{\tau - \hat{t}_2}{\tau}\right) \tag{Eq S77}$$

$$= s^{(m)} + g(T_f)\left(\frac{c_m}{c_1}\mu - \mu_m\right)\left(1 - \frac{f_2(R_2^{(c)})}{f_2(R_{2,\mathrm{in}})}\right). \tag{Eq S78}$$

Notably, in this limit, selection in coculture can remain distinct from selection in monoculture if the final toxin levels are sufficient to significantly decrease growth and the toxin is present for a significant fraction of the batch (equivalently, if the growth rate of the producer species at the input resource concentration is much larger than at the chemostat concentration). The expressions for the selection under the different conditions considered here are summarized in Table S2 .

**Toxin levels**

Note that the toxin level at the end of a batch ($T_f$) and at the chemostat steady-state ($T^{(c)}$) still appear in the expressions for selection. In order to get an expression for these quantities in terms of the model's parameters, we need to specify how the toxin is produced. One reasonable choice is to have the toxin production be proportional to the producer's growth rate. That is, the toxin dynamics within a batch are given by:

$$\dot{T} = p\dot{N_2} = pc_2 f_2(R_2)N_2. \tag{Eq S79}$$

In this case the toxin levels that determine selection become:

$$T_f = \int_0^\tau \dot{T}dt = \int_0^\tau p\dot{N_2}dt = p\left[N_2(\tau)) - N_2(0)\right] = pR_{2,in} \tag{Eq S80}$$

$$T^{(c)} = \frac{1}{d} pc_2 f_2(R_2^{(c)})N_2^{(c)} = p(R_{2,in} - R_2^{(c)}) = T_f - pR_2^{(c)}. \tag{Eq S81}$$

Therefore, in the boom-and-bust limit, the toxin levels that affect selection are larger than in the chemostat, but these toxin levels are only present during part of the time in each cycle. The difference between the toxin levels at the end of the cycle and the chemostat ($T_f - T^{(c)}$) is tightly linked to the growth rate of species 2; the faster species 2 grows, the smaller the difference. Replacing $R_2^{(c)}$ in Eq S81, we get:

$$T_f - T^{(c)} = pf^{-1}(\frac{\overline{d}}{c_2}) \tag{Eq S82}$$

or for a Monod-type consumption, as also visualized in Fig. S4:

$$T_f - T^{(c)} = p\frac{dK_2}{c_2 - \overline{d}} \tag{Eq S83}$$

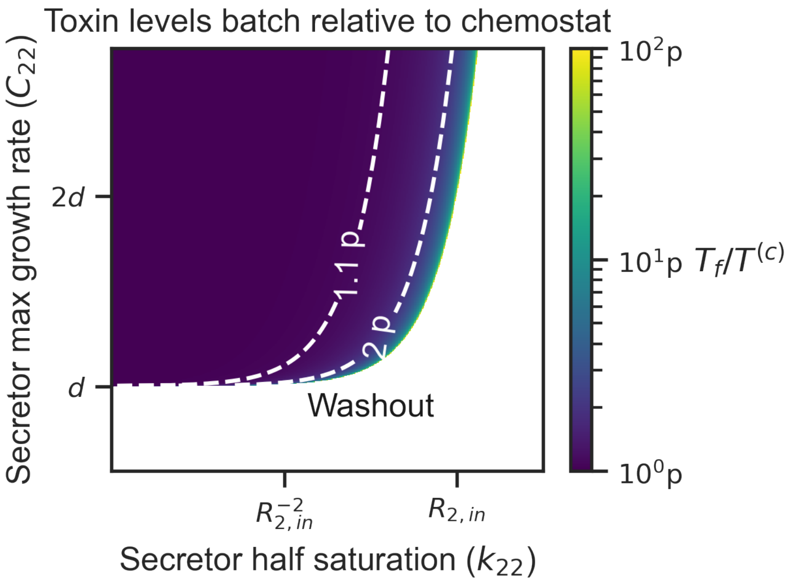

**Figure S4: Maximum toxin levels in dilution growth cycles are significantly different that those in chemostat only near washout.** the color denotes the log difference between $T_f$ and $T^{(c)}$, given the secretor maximum growth rate ($C_{22}$), and half-saturation constant ($k_{22}$). Maximum growth rate is given in units of the flow rate ($\bar{d}$), and $k$ is given in units of the provided resource concentration ($R_{2,in}$). white area denotes the area in which the secretor growth rate is slower than the flow rate and therefore it is washed out.

#### S6.2 Bacteriostatic toxin

We now consider a bacteriostatic toxin, which slows down the growth of the focal species rather than killing it. The within-batch dynamics are identical to the case of a bactericidal toxin except for the following:

$$\frac{1}{N_1}\dot{N}_1 = c_1 f_1(R_1)[1 - \mu g(T)] \quad \text{(Eq S84)}$$

$$\dot{R}_1 = -c_1 f_1(R_1) N_1 [1 - \mu_R g_R(T)] \quad \text{(Eq S85)}$$

$$\text{(Eq S86)}$$

That is, in addition to adapting the model such that the toxin slows down, rather than kill the focal species, we have also changed the resource consumption by species 1. This reflects the fact that a bacteriostatic toxin can also affect the yield. In particular, it may require more resources per cell. Note that, the monoculture situation, as well as the dynamics of $N_2$, $R_2$, and $T$ are identical to that of the bactericidal toxin case discussed above.

Selection for mutants in the maximal growth rate ($c_1$) and the maximal death rate ($\mu$) is given by:

$$s = \langle \gamma_m - \gamma \rangle = (c_m - c_1)\langle f_1(R_1)\rangle - (c_m\mu_m - c_1\mu)\langle f_1(R_1) g(T)\rangle \quad \text{(Eq S87)}$$

##### Chemostat limit

In coculture, here is again more resource at steady-state:

$$f_1(R_1^{(c)}) = \frac{\bar{d}}{c_1}\frac{1}{1 - \mu g(T^{(c)})} = f_1(R_1^{(m)})\frac{1}{1 - \mu g(T^{(c)})}, \quad \text{(Eq S88)}$$

and selection in coculture is given by:

$$s^{(c)}_{D\simeq 1} = \frac{d}{c_1}\frac{1}{1-\mu g(T^{(c)})}\left[(c_m - c_1) + (c_m\mu_m - c\mu)g(T^{(c)})\right] \quad \text{(Eq S89)}$$

$$= \frac{1}{1-\mu g(T^{(c)})}\left[s^{(m)} + \overline{d}\left(\frac{c_m}{c1}\mu_m - \mu\right)g(T^{(c)})\right]. \quad \text{(Eq S90)}$$

Note that these results are independent of the effect of the toxin on yield.

**Boom-and-bust limit ($D \gg 1$)**

As in the bactericidal case, we need to compute the approximate times in which each species grows ($\hat{t}_i$). If the focal species grows before the toxin production ($\hat{t}_1 < \hat{t}_2$), the toxin has no effect and selection is identical to the monoculture case. In the case where the toxin is produced before the focal species grows ($\hat{t}_1 > \hat{t}_2$), selection in this limit can be approximated as:

$$s^{(c)}_{D\gg 1} \approx \frac{\hat{t}_1}{\tau}(c_m - c_1)f_1(R_{1,in}) - \frac{\hat{t}_1 - \hat{t}_2}{\tau}(c_m\mu_m - c_1\mu)f_1(R_{1,in})g(T_f) \quad \text{(Eq S91)}$$

To find the growth time of the focal species, consider its overall growth over a batch

$$N_1(\tau) = N_1(0)e^{c_1 f_1(R_{1,\text{in}})\hat{t}_2 + c_1 f_1(R_{1,\text{in}})\left[1-\mu g(T_f)\right](\hat{t}_1-\hat{t}_2)} = N_1(0)e^{c_1 f_1(R_{1,\text{in}})\left[\hat{t}_1 - \mu g(T_f)(\hat{t}_1-\hat{t}_2)\right]}. \quad \text{(Eq S92)}$$

As before, we can use the fact that the focal species grows by a factor $D$ over the entire batch to get the growth time $\hat{t}_1$:

$$\log D = \log\frac{N_1(\tau)}{N_1(0)} = c_1 f_1(R_{1,\text{in}})\left[\hat{t}_1 - \mu g(T_f)(\hat{t}_1 - \hat{t}_2)\right], \quad \text{(Eq S93)}$$

giving

$$\hat{t}_1 = \frac{1}{1-\mu g(T_f)}\left[\frac{1}{c_1 f_1(R_{1,\text{in}})}\log D + \mu g(T_f)\hat{t}_2\right] = \frac{\log D}{c_1 f_1(R_{1,\text{in}})(1-\mu g(T_f))}\left[1 + \frac{c_1 f_1(R_{1,\text{in}})}{c_2 f_2(R_{2,\text{in}})}\mu g(T_f)\right]. \quad \text{(Eq S94)}$$

The growth times we derived can be plugged into the expression for the selection to get a final expression that depends only on the model parameters. However, more insight can be gleaned by relating $\hat{t}_1$ to the difference between the growth times:

$$\hat{t}_1 = \frac{1}{c_1 f_1(R_{1,\text{in}})}\log D + \mu g(T_f)(\hat{t}_1 - \hat{t}_2). \quad \text{(Eq S95)}$$

We can now also express the selection as a function of the difference between the growth times:

$$s^{(c)}_{D\gg 1} \approx s^{(m)} + c_m(\mu - \mu_m)f_1(R_{1,in})g(T_f)\frac{\hat{t}_1 - \hat{t}_2}{\tau}. \quad \text{(Eq S96)}$$

This shows that selection in the large dilution limit does not necessary converge to the selection in monoculture. It can be distinct from it if there is a non-negligible fraction of the cycle where the focal species is growing and the toxin is present. That is, if the toxin is produced early in the growth of the focal species.

Notably, if mutations only affect maximal growth rate ($C_1$) and not maximal toxin inhibition ($\mu$), selection in coculture is also equal to selection in monoculture.

**Summary**

The expressions for the selection under the different conditions considered here are summarized in Table S2 .

| | Chemostat | Large D |
|---|---|---|
| Monoculture | $\frac{c_m - c_1}{c_1}\overline{d}$ | $\frac{c_m - c_1}{c_1}\overline{d}$ |
| Coculture cidal | $s^{(m)} + \left[\frac{c_m}{c1}\mu - \mu_m\right] g(T^{(c)})$ | $s^{(m)} + \left[\frac{c_m}{c_1}\mu - \mu_m\right] g(T_f)\left(\frac{\tau - \hat{t}_2}{\tau}\right)$ |
| Coculture static | $\frac{1}{1-\mu g(T^{(c)})}\left[s^{(m)} + \overline{d}\left(\frac{c_m}{c1}\mu_m - \mu\right) g(T^{(c)})\right]$ | $s^{(m)} + c_m(\mu - \mu_m) f_1(R_{1,in}) g(T_f)\frac{\hat{t}_1 - \hat{t}_2}{\tau}$ |

Table S2 : Selection with a toxin

#### S7 Simulating toxin-mediated interactions

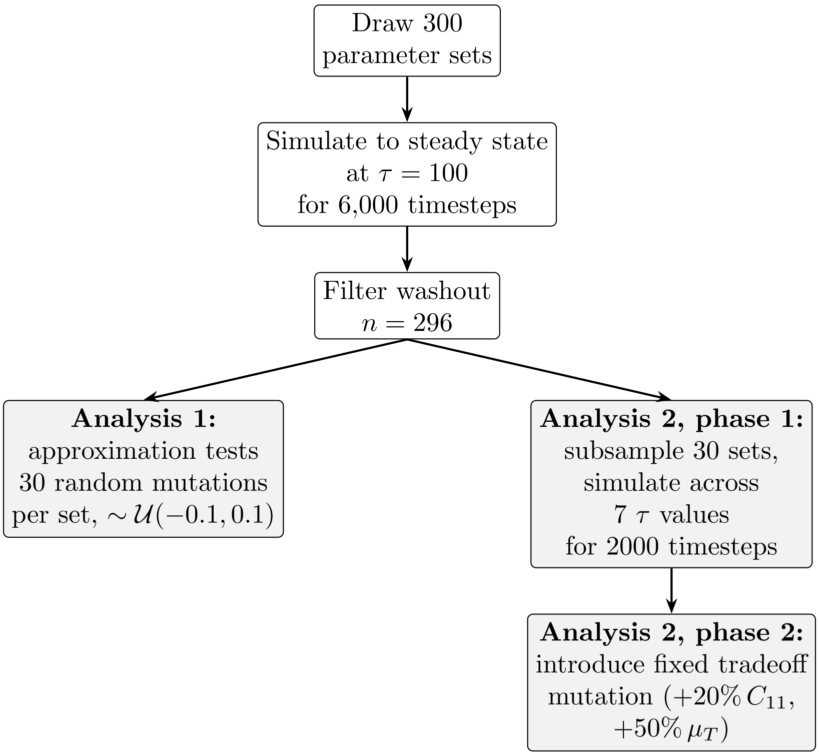

**Figure S5:** Simulation pipeline for toxin-mediated interactions.

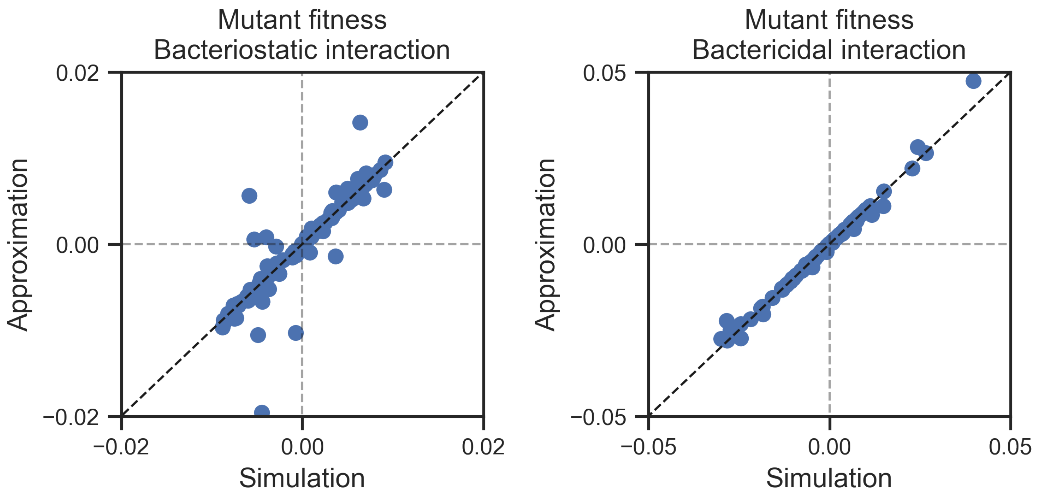

**Figure S6: Approximations accurately reproduce mutant fitness in simulated toxin secretion pairs.**

Table S3 : Parameter distributions and simulation counts for toxin-mediated interactions.

| **Parameter** | **Distribution / Value** | **Notes** |
|---|---|---|
| *Species growth parameters* | | |
| $c_{11}$ (focal max growth rate) | Uniform$[2, 4]$ | Focal consumes $R_1$ only |
| $c_{22}$ (secretor max growth rate) | Uniform$[0.2, 3]$ | Secretor consumes $R_2$ only |
| $k_{11}, k_{22}$ (half-saturation) | Log-uniform on $[10^6, 10^8]$ | 50 discrete values |
| *Toxin parameters* | | |
| $\mu_T$ (max toxin effect) | Log-uniform on $[10^{-2}, 10^0]$ | |
| $k_T$ (toxin half-effect conc.) | Log-uniform on $[10^4, 10^7]$ | |
| $h$ (Hill coefficient) | Uniform$[1, 7]$ | |
| $p$ (toxin production rate) | 1 | Fixed |
| *Environmental parameters* | | |
| $R_{1,\text{in}}$ (focal supply) | $10^9$ | Fixed |
| $R_{2,\text{in}}$ (secretor supply) | Log-uniform on $[10^7, 10^8]$ | |
| *Growth regime* | | |
| $\overline{d}$ (avg. dilution rate) | $\log(D)/\tau$ | Set per parameter set |
| $D$ (dilution factor) | $10^4$ (Analysis 1); $\exp(d\tau)$ (Analysis 2) | |
| $\tau$ (Analysis 1) | 100 | Fixed |
| $\tau$ (Analysis 2) | $\{5, 10, 50, 100, 200\}$ | 5 values |
| *Mutations* | | |
| Mutant introduction abundance | $10^{-5}$ | Both analyses |
| Analysis 1 mutations | $\Delta c_{11}, \Delta\mu_T \sim \mathcal{U}(-0.1, 0.1)$ | 30 per set |
| Analysis 2 mutation | $\Delta c_{11} = +20\%$, $\Delta\mu_T = +50\%$ | Fixed tradeoff |
| *Simulation counts* | | |
| Parameter sets drawn | 300 | |
| Sets passing washout filter | 296 | |
| Analysis 1 total mutant invasions | $296 \times 30$ | At $\tau = 100$ |
| Analysis 2 total mutant invasions | $30 \times 5$ | Across $\tau$ |
